# Proximal sesamoid bone fatigue in Thoroughbred racehorses is associated with palmar osteochondral disease in the adjacent third metacarpal bone condyle

**DOI:** 10.64898/2026.10.20.760467

**Authors:** Elleana J. Thom, Soroush Irandoust, Corinne R. Henak, Peter Muir

**Author notes:** Correspondence: Dr. Peter Muir, Dr. Soroush Irandoust. These authors contributed equally to the study.

## Abstract

**Background:** Proximal sesamoid bone (PSB) stress fracture is a common catastrophic injury in racing Thoroughbreds that is associated with bone sclerosis and development of focal osteolysis. Standing computed tomography (sCT) enables sensitive identification of these structural changes and associations with other more common fetlock fatigue injury such as palmar/plantar osteochondral disease lesions affecting the condyles of the third metacarpal/metatarsal (MC3/MT3).

**Objective:** To gain insight into the PSB stress fracture mechanism, we examined associations between fatigue injury prevalence in the three major metacarpophalangeal (MCP) sites: PSB, parasagittal grooves (PSG), and third metacarpal bone MC3 condyles.

**Study design:** Retrospective clinical study.

**Methods:** We studied 22 catastrophically injured Thoroughbred racehorses. Thoracic limbs were examined using computed tomography (CT) (n=29). Subjective CT assessment classified 58 PSBs as 32 control and 26 case PSBs with bone fatigue (transverse fracture, focal osteolysis, or severe sclerosis). Opposing joint surfaces were also studied. PSB bone mineral density (BMD) was estimated.

**Results:** Prevalence of fatigue injury in the PSB, condyle, and PSG was 44.8% (n=26), 43.1% (n=25), and 17.2% (n=10), respectively. For MC3 condyles with POD, 88.0% had fatigue injury in the corresponding PSB (OR = 53.2). Prevalence of PSB fatigue injury was 20.0% when there was PSG fatigue injury. The odds of a PSB having fatigue injury when the opposing MC3 PSG had signs of fatigue injury was 0.25. PSBs with opposing MC3 condyles containing a POD lesion had significantly increased BMD compared with opposing control condyles. BMD was significantly increased in PSBs with structural change associated with cyclic fatigue.

**Conclusions:** Development of PSB fatigue and adaptive PSB sclerosis is associated with POD. Stress fractures in Thoroughbreds commonly affect the MCP joint. Condylar fractures occur from PSG or POD fatigue injury. PSB fractures are also common. Our findings suggest detailed clinical assessment of PSB structural change in racing Thoroughbreds diagnosed with POD is merited as part of routine injury prevention screening.

## 1 INTRODUCTION

Musculoskeletal injuries are the most common cause of death in Thoroughbred racehorses, many of which are stress fractures in the metacarpophalangeal (MCP) joint. These fatigue-induced fractures can occur in the condyles of the third metacarpal bone (MC3) from fatigue injury in the parasagittal groove (PSG) or a palmar osteochondral disease (POD) lesion, or in the proximal sesamoid bones (PSB), with the latter being the most common MCP stress fracture in the United States and Hong Kong in catastrophically injured horses.^1,2^ Functional outcome after surgical treatment of serious PSB stress fractures is often poor and in one study only 28% of horses raced again after surgery.^3^ PSB stress fractures may affect one or both bones within a fetlock. Smaller avulsion fractures are commonly associated with lameness in racehorses and can be successfully treated.^4,5^ Complete comminuted or transverse mid-body fracture affecting both PSBs with loss of fetlock support represent a serious career changing injury that often leads to euthanasia.^6–8^

Risk factors for catastrophic suspensory apparatus injury (including PSB injury) in Thoroughbred racehorses include increasing age,^9^ male sex,^9^ horseshoe toe grabs,^10^ horseshoe pads,^11^ increasing hoof toe-heel angle,^12^ evidence of mild suspensory apparatus injury found during physical examination,^13^ actively training but not racing,^9^ greater time between layups, 2 to 5 career races, and high exercise intensity.^11^ Due to the anatomy of the fetlock, there is no reliable clinical approach for detecting horses with a heightened risk of PSB stress fracture. PSB bones with concerning structural change cannot be reliably identified using planar digital radiography (DR)^14^ but are much more readily detected using computed tomography (CT) imaging.^15,16^ Positron emission tomography (PET) has also been used to examine PSB metabolic activity associated with bone remodeling in Thoroughbred racehorses and obtain spatial identification of the location of remodeling,^17^ which may help with identification of PSBs at risk of stress fracture.

Very high cyclic compressive loads are transferred from the PSB bones to the condyles of the distal end of the MC3/MT3 bone during training and racing. Peak load and contact stress on the palmar third metacarpal condyles are estimated at ∼26kN and 61MPa, respectively.^18^ In response to repetitive high cyclic fatigue loading, adaptation occurs in the form of sclerosis in both the MC3 condyles as well as the PSBs.^19^

Bone is normally a self-repairing material that is renewed through the process of bone remodeling. A proportion of remodeling is targeted to the repair of fatigue microdamage.^20^ If microdamage accumulates in the face of reparative remodeling, bone structural properties are degraded^21,22^ and subchondral osteolysis may form in the MC3 condyles (POD), focally in the PSG.^23^ Focal osteolysis may also form in the subchondral or palmar regions in the PSB (**Figure 1**),^24^ all of which may elevate risk of a complete stress fracture of the PSB in isolation or combination with condylar stress fracture. Parasagittal condylar stress fracture arises from an accumulation of fatigue microdamage in subchondral bone.^25,26^ Similarly, PSB fatigue in the subchondral axial/abaxial or palmar/plantar bone leads to focal sclerosis, osteopenia, development of an incomplete articular fracture line, and osteoarthritis.^2,6,19,24,27^ Such structural changes are potentially associated with elevated risk of serious or fatal stress fracture.^2,6,15,18,24,27^

**Figure 1.**
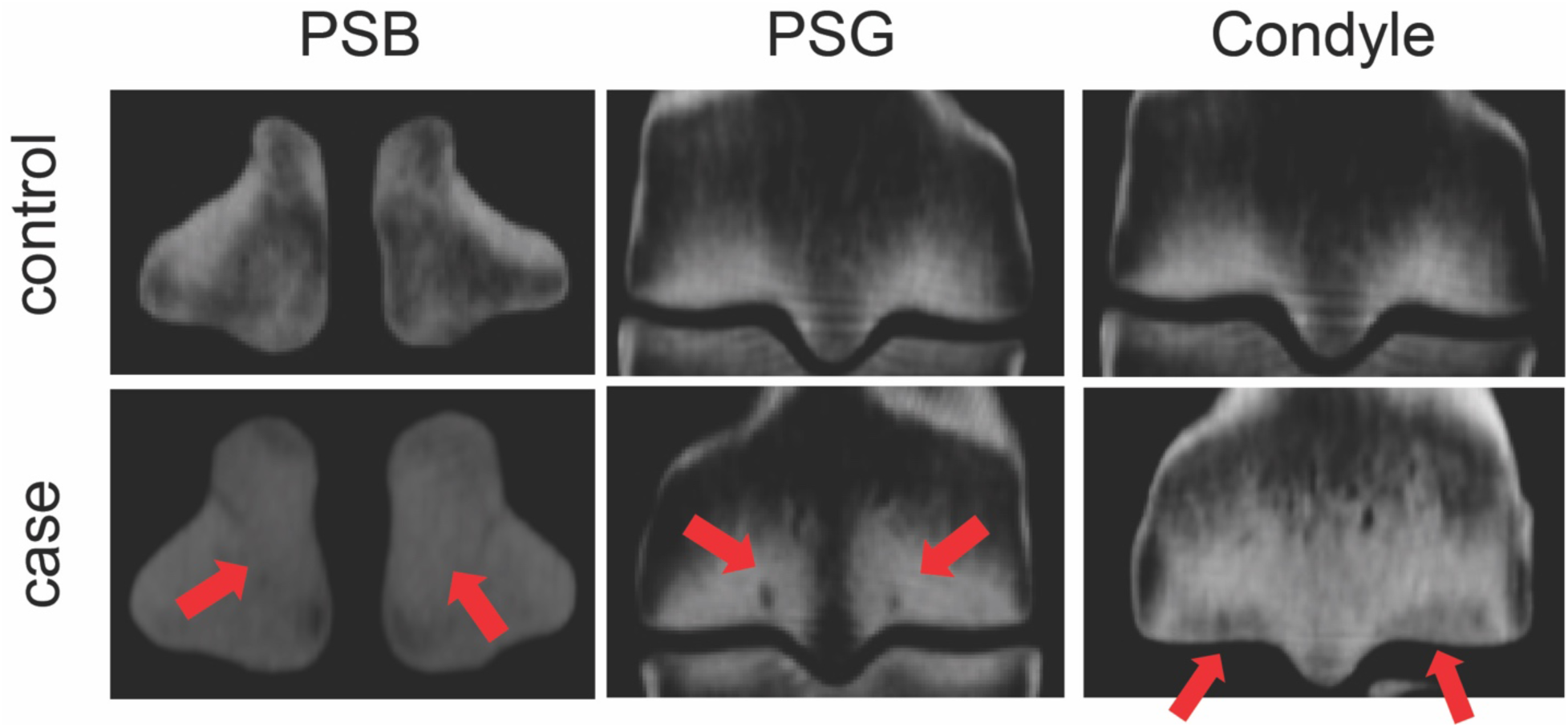
Frontal sCT reconstructions of the proximal sesamoid bones (PSB), parasagittal grooves (PSG), and third metacarpal (MC3) condyles, with and without structural changes associated with bone fatigue. The case PSB shown is sclerotic as shown with the arrows with associated enlarged vascular channels. The case PSG shown has biaxial focal osteolysis indicated with the red arrows. Palmar osteochondral disease (POD) in the MC3 condyles is seen as subchondral lucency (red arrows).

Identification of signs of fatigue injury and adaptive and reparative responses are challenging with DR, especially in the PSBs, but can be better achieved with standing CT imaging (sCT).^28^ Currently, the typical location for stress fracture initiation is not understood.^29^ Histological studies of bone microdamage have not consistently shown increased PSB microdamage in racehorses with fractures.^29–31^ In contrast, a microCT study found sclerosis, microdamage accumulation, and focal areas of lysis in racehorses with cumulative racing, observations consistent with bone fatigue in the PSB leading to fracture.^2^ These inconsistent observations are likely a consequence of study design, cyclic strain levels, the burden of microdamage, duration of loading, and stressed volume effects^32^ during fracture initiation. Taken together, current evidence indicates that transverse mid-body and basilar PSB fractures in Thoroughbred racehorses are stress fractures with a similar mechanism to condylar stress fractures where fracture propagation is often associated with development of focal lucency in the PSB subchondral plate or flexor cortex.^27,31^

To gain more insight into the PSB stress fracture mechanism, the goal of this retrospective study was to use CT imaging to examine associations between prevalence of bone fatigue in the three major sites in the MCP joint: PSBs, PSGs, and MC3 condyles *ex vivo*, as potential relationships between development of bone fatigue on these opposing joint surfaces are not understood. We hypothesized that PSB fatigue is associated with POD of the opposing MC3 condyle in catastrophically injured horses. We additionally hypothesized that MC3 condyles with POD had opposing PSBs with significantly higher BMD from sCT imaging and segmentation of the PSB volume for density estimation. Confirmation of these hypotheses could lead to beneficial improved detection of PSB fatigue injury clinically through sCT fetlock imaging of horses found to have POD using more routine DR imaging.

## 2 Methods

Because PSB stress fracture is most common in the thoracic limb, *ex vivo* imaging in this study was limited to the thoracic limb. This helped generate a standardized image set for evaluation of structural changes.

### 2.1 Limbs and PSBs

Entire distal thoracic limb specimens were obtained from 22 Thoroughbred racehorses that died or were euthanatized for reasons unrelated to the present study during athletic training and racing in Florida, USA, and were given for use in this work. Horses were euthanatized humanely by a veterinarian at the racetrack because of catastrophic injury. Limbs were transected at the level of the carpus, sealed in plastic bags, and stored at −20°C until needed.

### 2.2 Fetlock computed tomography

Each limb specimen was scanned frozen to enable specimen preservation and to minimize tissue exposure to freeze-thaw cycles.^23^ Scanning frozen does not alter CT imaging of osseous structures.^33^ CT imaging was performed at high-resolution using 0.55mm contiguous slices using a robotic CT scanning system, which has the capability of performing vertical scanning of limb pairs as well as horizontal scanning of the head/neck in the standing sedated horse. The 24-slice helical Equina^®^ scanner (Asto CT Equina^®^) is a fixed installation and has a 240V single phase 30A uninterruptable power supply. Scanning was performed using an exposure of 160kVp and 8mA at 1 second per 360° revolution using 24 detector rows with a variable helical pitch, typically 0.55. Slice acquisition rate is 36 slices/sec with an image acquisition matrix of 1024×1024 and a resolution at isocenter of 0.75mm. Maximum scanning distance is 100cm vertically at 2cm/second.^16^ No scout images are needed. Axial scanning of limbs was performed in the horizontal scanning mode with the isolated limbs placed on the positioning table normally used for head scanning in the standing horse. DICOM images were then stored on a network drive. CT images of a subset of 29 limbs from 22 horses (7 paired-limb scans and 15 single-limb scans) were used for assessment of fatigue damage in the PSBs, PSGs, and MC3 condyles. Work flow is summarized in Figure S1. One fetlock with two comminuted PSB fracture were excluded from limb pairs from one horses because the severity of comminution made classification as a PSB stress fracture difficult and impaired accurate segmentation (see below), and a single limb from this horse was included in the analysis. The medial and lateral aspects were both assessed, creating a sample size of n=58 PSBs representing 29 limbs from 22 horses.

An electron density phantom (model 062M, CIRS Inc, Arlington, VA) with 4 calcium hydroxyapatite (HA) plugs with densities of 200, 800, 1250, and 1750 mg/cm^3^ was also scanned with the same scanner at the same exposure settings to find the equivalent radiological or CT density, *ρ_CT_* (*mg_HA_*/*cm*^3^), from the CT number in Hounsfield units (HU):

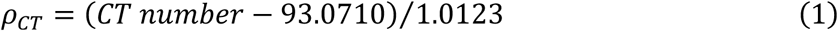

### 2.3 Computed tomography image evaluation

Each joint region (PSG, PSB, Condyle) was classified as a control or a case region with signs of bone fatigue using an experienced observer (PM). Cyclic fatigue of the PSB included mid-body or basilar transverse stress fracture of one or both PSBs within a fetlock, subchondral/palmar lucency, and moderate to extensive sclerosis.^2,24,27^ PSG bone fatigue included condylar fracture through the PSG, and subchondral focal osteolysis in this region.^14^ Bone fatigue in the MC3 condyles included those with focal subchondral osteolysis with surrounding sclerosis (POD) with or without condylar stress fracture.^28,34^ Bone fatigue features were graded as present or absent.

### 2.4 Segmentation of the proximal sesamoid bones

Mimics (v.26) was used to segment the PSB. Using a gradient-based segmentation tool, the external surface of the bone was detected semi-automatically for all individual sCT image slices sequentially. All reconstructable fragments of the fractured PSBs were segmented separately and included in the same mask for further analysis. Using morphology operations, two pixels was eroded from the external surface to ensure its alignment with the bone surface, and then the 3D PSB was constructed and smoothed. The pixel intensities of each PSB in HU was then exported for further data analysis.

### 2.5 Data analysis

First, prevalence of fatigue injury in each of six regions of the MCP joint (lateral PSB, lateral condyle, lateral PSG and equivalent medial regions) was calculated. Next, contingency tables were created to find associations between these injuries by calculating relative risk and odds ratio. Finally, HA-equivalent BMD and its distribution (using kernel density estimation) were calculated for all PSBs and probability density plots were generated. We also compared PSB BMD between those with opposing MC3 condyles with and without POD. A linear mixed model (LMM) was made with PSB BMD as the dependent variable, and independent variables as follows: POD status and PSB fracture status as fixed-effect categorical variables, limb and condyle (right/left, medial/lateral) sides as fixed-effect categorical covariates, and horse-ID with random effect. Another LMM was made only to compare BMD between case and control PSBs, with PSB case/control status as the fixed categorical variable, limb and condyle (right/left, medial/lateral) sides as fixed-effect categorical covariates, and horse-ID with random effect.

Results were considered significant at *p*<0.05. The average PSB BMD distributions with 95% confidence intervals were plotted for visual comparison. Data analysis was performed in Python 3.12.3 with numpy 1.26.4, pandas 2.2.0, scipy 1.13.1, scikit-learn 1.5.2, statsmodel 0.14.0, and matplotlib 3.9.2.

## 3 Results

Subjective clinical assessment classified the 58 joint regions studied as 22 control regions and 36 case regions with clinically important structural change, based on review of CT images (**Figure 2**). Of the case PSBs, 12 had fractures. PSB fractures were biaxial in 5 joints. Both mid-body transverse fractures with focal subchondral lysis and sclerosis affected the medial PSB with associated lateral mid-body PSB transverse fracture in a sclerotic bone and collapse of the fetlock (**Table S1**). Prevalence of bone fatigue in the PSB, condyle, and PSG was 44.8% (n=26), 43.1% (n=25), and 17.2% (n=10), respectively (**Figure 2**). Of the MC3 condyles with POD, 88.0% had structural change associated with bone fatigue in the corresponding PSB. The odds of PSB fatigue if POD was detected in the opposing MC3 condyle was 53.16 (*p* < 0.0001) (**Table S2**). Case PSBs had significantly higher BMD than control PSBs (*p*<0.001, **Figure S2A**). The BMD profile plot of case PSBs were left-skewed with higher proportion of bone tissue with 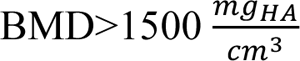 compared with control PSBs (**Figure S2B**). Fractured PSBs showed qualitatively higher BMD than non-fractured PSBs (**Figure 2CD**), although statistical significance measurement was not possible due to the small number of horses with PSB fractures. Nine of the case PSGs had focal subchondral lysis. Three of the case PSGs had a condylar fracture. One case condyle had a POD-associated condylar fracture. The prevalence of having PSB fatigue was only 20.0% when there was PSG fatigue injury. The odds of PSB fatigue when the opposing MC3 PSG had signs of fatigue injury was 0.25 (**Table S3**).

**Figure 2.**
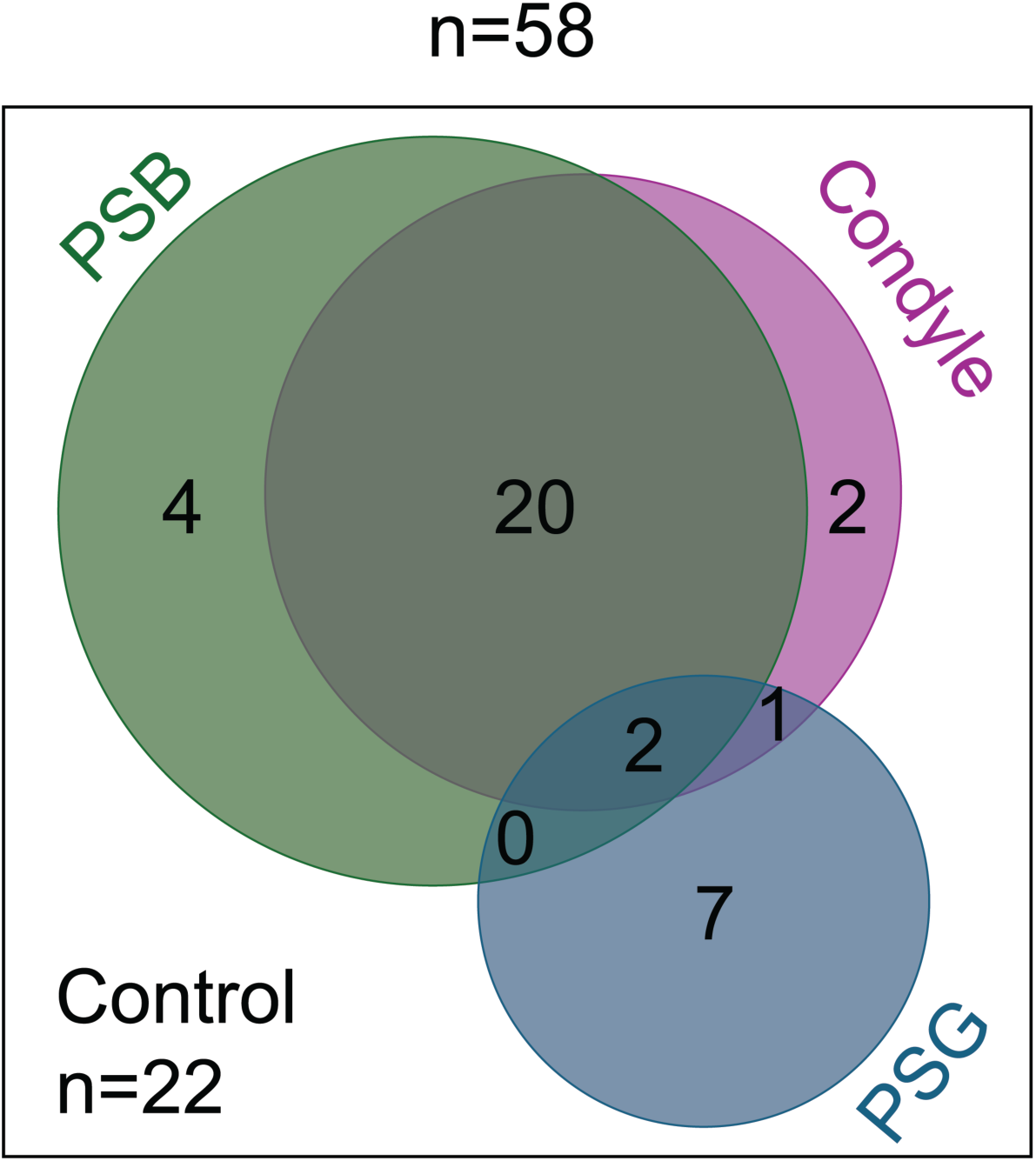
The Venn diagram shows extensive overlap between having palmar osteochondral disease (POD) in the condyle and structural change associated with fatigue in the proximal sesamoid bones (PSBs). Focal osteolysis or fracture in the parasagittal groove (PSG) of the third metacarpal is an almost completely isolated change from development of POD in the condyle or PSB fatigue.

Additionally, our analysis revealed a minimal correlation between the presence of PSG lysis/fracture and POD in the condyles. Specifically, the odds of PSG fatigue injury in MC3 condyles with POD was 0.51 (**Table S4**). The only fetlock joint region with both POD and PSG lysis had highly sclerotic PSB as well.

PSBs with opposing condyles with POD (**Figure 3**) had increased BMD (*p*<0.001, **Figure 4A**) than those with an opposing control MC3 condyle. The BMD profile plot of these PSBs were left-skewed with higher proportion of bone tissue with BMD>1500 *mg_HA_*/*cm*^3^ than POD controls (**Figure 4B**). We also found that PSBs with fractures had qualitatively higher BMD compared with those with non-fractured bones (*P*=0.5). However, due to the small number of horses with PSB fracture (n=6, **Figure 5**), the statistical analysis was underpowered (**Figure S2**) for significance calculations.

**Figure 3.**
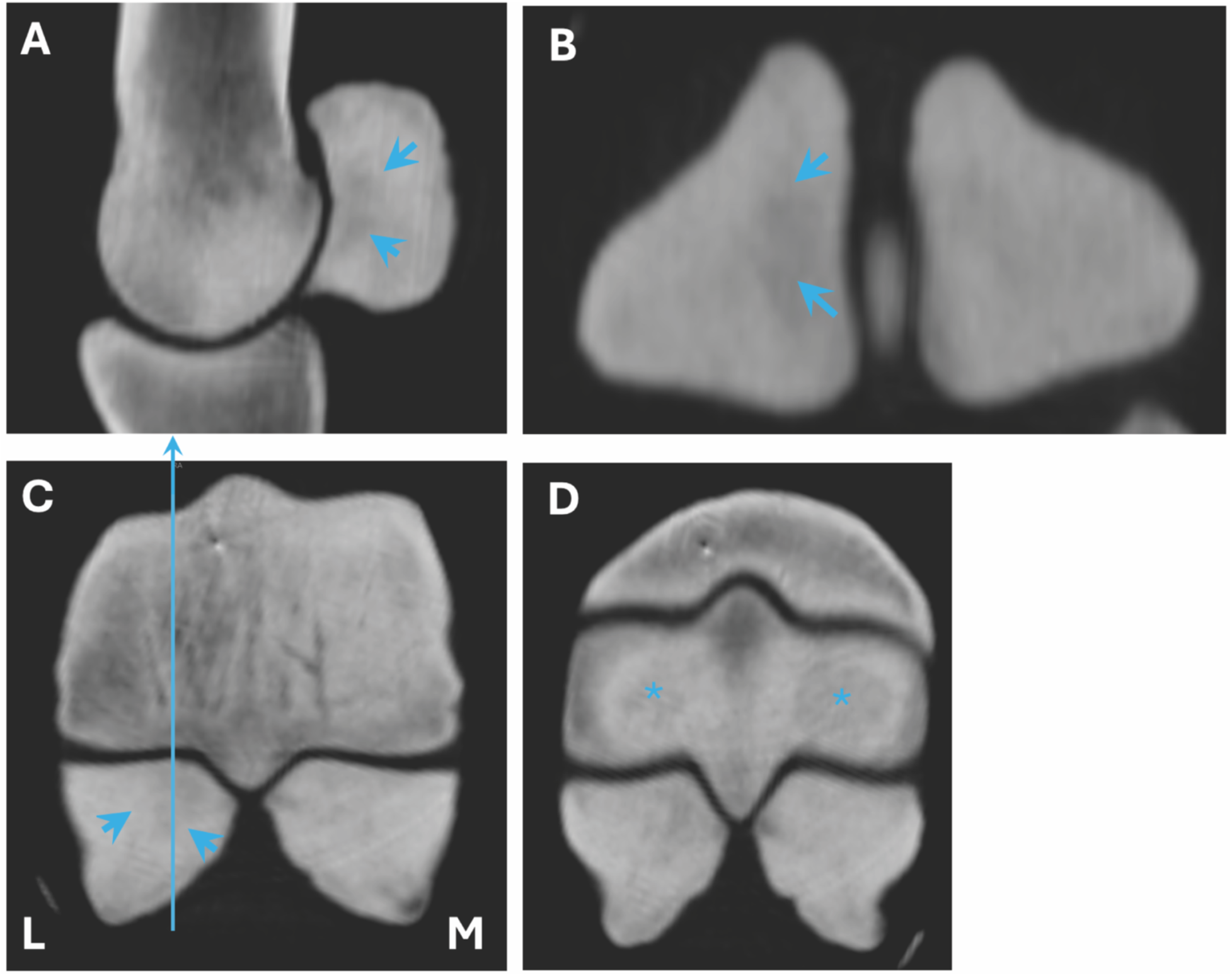
Sagittal (**A**), frontal (**B**), transverse (**C**), and dorso45proximo-palmarodistal oblique (**D**) postmortem computed tomography images of the right fetlock of a Thoroughbred racehorse with catastrophic injury. L – lateral, M – medial. There are large palmar osteochondral disease (POD) lesions in both condyles (*). Sclerosis is both proximal sesamoid bones is present together with axial focal subchondral lysis (arrowheads) in the lateral bone because of fatigue injury.

**Figure 4.**
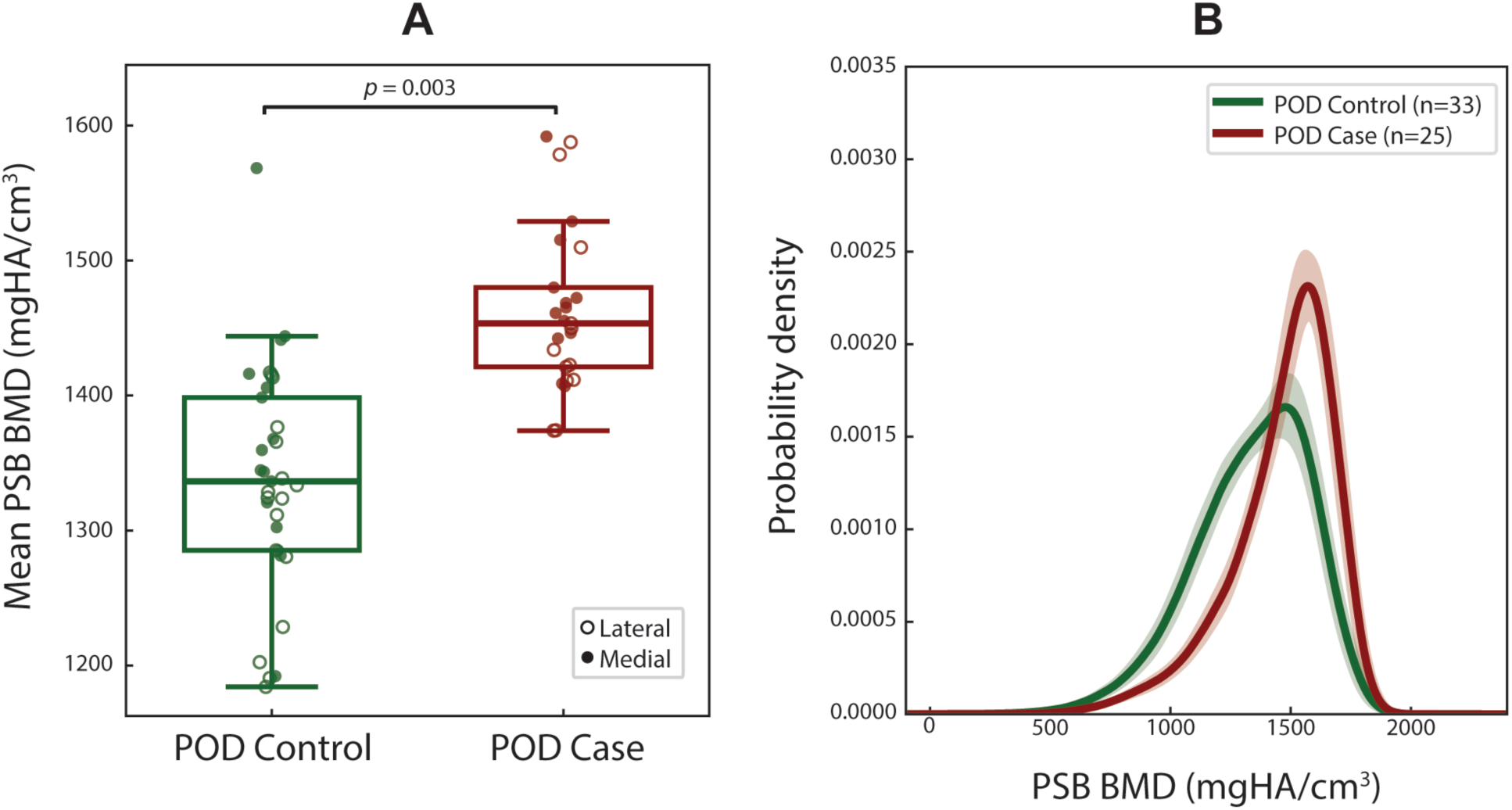
Proximal sesamoid bones with POD in the opposing MC3 condyle had significantly higher average HA-equivalent density (A) with a left-skewed distribution (B). Shaded areas in the profile plot indicate the 95% confidence interval.

**Figure 5.**
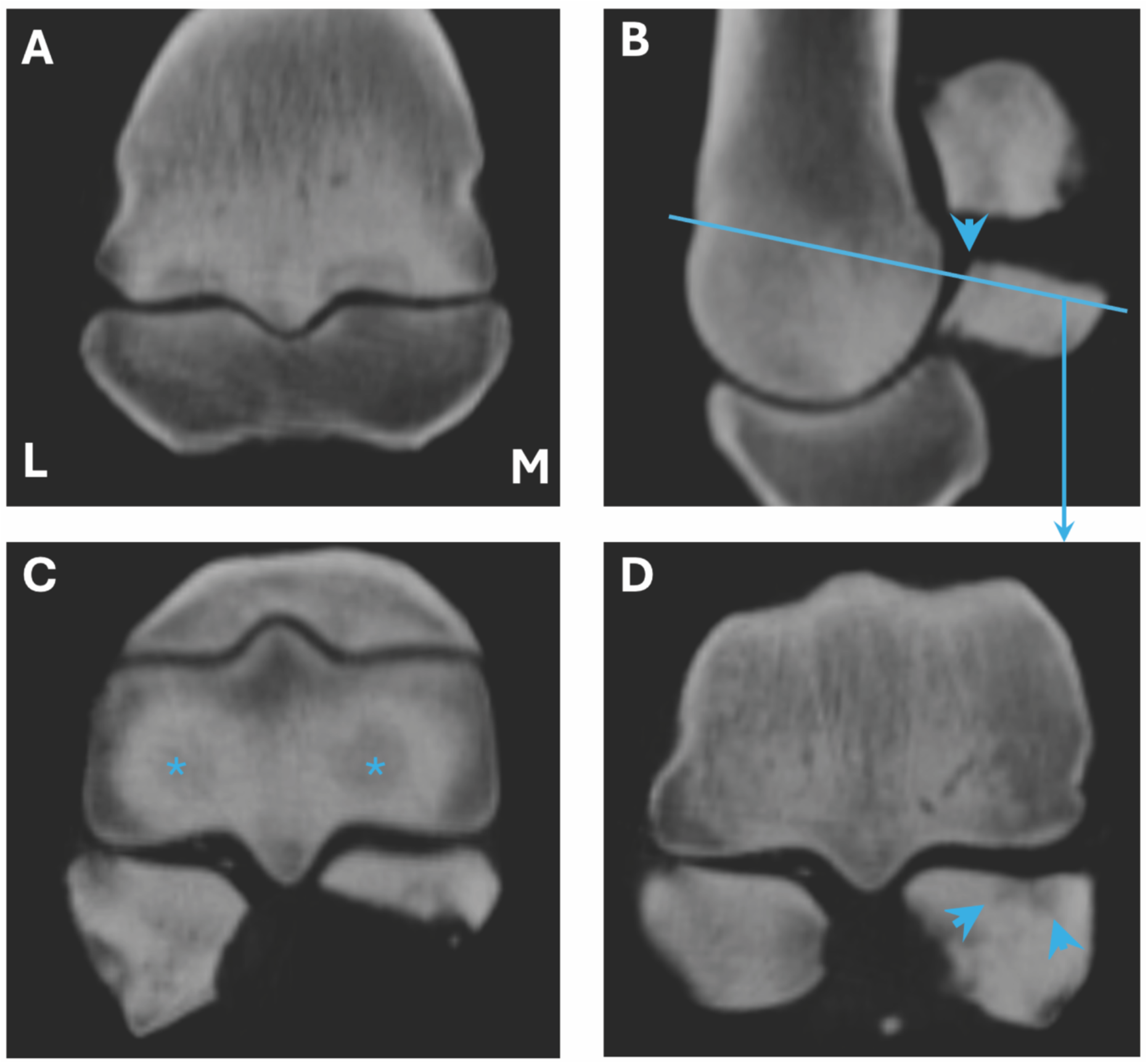
Dorso45proximal-palmarodistal oblique frontal (**A**), sagittal (**B**), and transverse (**C, D**) postmortem computed tomography images of the left fetlock of a Thoroughbred racehorse with catastrophic injury. L – lateral, M – medial. There are large palmar osteochondral disease lesions in both condyles (*). Sclerosis in both proximal sesmoid bones (PSB) is present together with a mid-body transverse stress fracture of the medial PSB with some minor comminution (not shown). The PSB stress fracture is associated with a large region of abaxial focal subchondral lysis (arrowheads) representing subchondral fatigue injury.

## 4 Discussion

Currently, PSB stress fracture is an important cause of Thoroughbred racehorse morbidity and death.^1,2^ In Thoroughbred racehorses, microdamage and associated subchondral sclerosis of the palmar/plantar condyles of the MC3/MT3 bone and the PSBs develops because of race training.^2,28,35^ In some horses, site-specific accumulation of fatigue damage eventually leads to development of a focal osteolytic lesion in the PSB in either the subchondral plate or the flexor cortex.^2,27^ In other horses, severe PSB sclerosis will develop as an adaptive response that includes modeling to increase local BMD as well as remodeling targeted to the repair of microdamage. Findings from the present study confirm our hypothesis that POD in the MC3 condyles and signs of cyclic fatigue and bone structural change in the PSBs were highly associated in catastrophically injury horses. This association is logical because if the subchondral plate of a joint surface is exposed to cyclic fatigue during athletic activity, the opposing joint surface would be expected to experience similar fatigue loading.

Cumulative athletic history of intensive joint loading contributes to accumulation of fatigue microdamage with career progression and the risk of distal limb fracture increases with increasing horse age.^2,35,36^ Independent of the age of the horse, microdamage levels are lower in resting horses, which is consistent with the observation that higher levels of subchondral bone remodeling develop during rest periods.^35^ Although microdamage cannot be detected in living horses with current imaging methods, the extent of radiographic subchondral bone sclerosis is correlated with microdamage^2,35,36^ which is readily identified by sCT imaging. In this regard, we were also able to confirm our second hypothesis that PSBs had significantly higher BMD when there is a POD lesion on the opposing condyle, suggesting PSB fatigue and associated bone injury is promoted by development of POD. At other stress fracture predilection sites in

Thoroughbred racehorses, there is typically a local reduction in material properties with development of sclerosis.^37^ Abrupt changes in BMD adjacent to focal osteolytic regions where microdamage is being remodeled, or areas of sclerosis, may facilitate initiation of a critical stress crack during fracture propagation.^20^ Our findings in a small number of cases of biaxial PSB fracture with collapse of the fetlock from loss of palmar support sugguest that development of PSB sclerosis and subchondral focal osteolysis is particularly concerning regarding risk of imminent serious or fatal injury. Effects of pathological change on PSB material properties warrants further investigation.

Although existence of obvious bone fatigue in PSG and PSB predilection sites is rare, such cases may have a significantly higher risk of stress fracture,^15,25,26,27,32^ in contrast to the condyle, where POD is relative common.^27^ A substantial proportion of condylar stress fractures are associated with POD, particularly in the USA,^38,39^ but many condylar fractures are associated with PSG subchondral lysis.^14,25,26,40^ The risk that POD lesions pose to development of PSB stress fracture is currently also a gap in knowledge. In the horses of the present study racing in the USA, PSB fatigue was commonly associated with development of POD, but this may not be true of Thoroughbred populations racing in other parts of the world. In horses racing in Australia, POD lesions were commonly found in a recent study, but PSB focal lucency and/or fracture was not.^28^ The potential relationships between POD lesion severity and PSB fatigue, PSG fatigue, and abaxial condylar stress fracture needs further investigation. Estimates of the potential impact of using sCT to assess horses with planar radiographic evidence of POD would be helpful. In this regard, our results suggest segmentation of the PSB volume in a sCT for estimation of PSB density seems like a promising objective continuous variable that reflects development of PSB fatigue through an adaptive bone sclerosis response to cyclic fatigue loading during training and racing.

Early detection of PSB fatigue as part of injury prevention screening remains an important research goal as severe fetlock injuries reduce athletic performance, lead to early retirement, and limit a horse’s future career in other pursuits.^41^ sCT imaging is a practical noninvasive diagnostic imaging tool that enables measurement of BMD and detection of PSB structural changes, in contrast to DR. Such information enables objective assessment of PSB fatigue damage. In absence of access to sCT imaging, higher likelihood of the presence of fatigue damage in the PSBs should be considered in cases with POD in the MC3 condyles, which are easier to identify with DR^42^ or sCT due to the larger lesion size. Given the expected high prevalence of POD lesions and the low prevalence of PSB fatigue in the general racehorse population based on the current literature, the positive predictive value of a POD lesion being associated with a PSB fatigue or bone injury may be low. True prevalence needs to be determined in future work, as PSB fatigue injury may be more common than expected because it is easy to miss clinically. The clinical consequences of failing to identify Thoroughbred racehorses with heightened risk of serious or fatal PSB stress fracture is high, so the linkage between POD and PSB fatigue injury is an important one.

There were several limitations to this study. All limb specimens were collected in the United States which could influence findings when it comes to the prevalence of the three types of MCP fatigue injury we studied compared to Thoroughbred racehorses from different parts of the world, as interactions between limb loading from athletic activity and racing surface properties, microdamage accumulation and targeted bone remodeling are complex. The limited clinical information regarding these horses also prevented more detailed epidemiological analysis of clinical factors associated with PSB fatigue injury. Subjective classification of condyle, PSG, and PSB bone with regarding to evidence of fatigue injury was performed by a single obsever to ensure consistent grading across the dataset. It is possible that other observers may have made different assessments. CT imaging of the isolated limbs were acquired in a different manner from sCT imaging clinically.^16^ However, CT imaging can be reconstructed easily in different planes to enable standard clinical evaluation irrespective of how the images were acquired.

## Conclusions

In the present study, we obtained new insight into associations between bone fatigue in the three major injury predilection sites in the MCP joint in Thoroughbred racehorses. We identified a strong association between development of subchondral fatigue injury in the MC3 condyle (POD) and the presence of PSB fatigue and bone injury with an odds ratio of 53.2. PSB stress fractures are common across the world where Thoroughbreds race and train and this injury is of public concern.^41^ Our findings are expected to impactfully improve objective clinical stress fracture risk assessment for injury prevention screening of Thoroughbred racehorses as sCT imaging becomes more widely used for clinical risk assessment, as detection of POD in the fetlock is easier than PSB structural change. Our findings suggest evaluation for structural changes in the PSBs is a higher priority in horses with POD lesions. More epidemiological study of these features in Thoroughbred populations undertaking race training is needed including any relationship with POD severity.

## Acknowledgements

The authors would like to thank Dr. David Ergun for his help with acquiring the sCT image sets. We are also grateful to Dr. Patricia Marquis who contributed limb specimens for analysis.

## Data Integrity Statement

Dr. Muir and Dr. Irandoust had full access to all of the data in the study and take responsibility for the integrity of the data and accuracy of the data analysis.

## Data Availability Statement

The clinical data used to support the findings of this study are included within the article.

## Ethical animal research

This is an *ex vivo* study. Research ethics committee oversight not required for an ex vivo study performed on material obtained from a disposal service.

## Informed Consent

Not applicable. Equine samples were sourced via a disposal service.

