## Supplementary Data for "Proximal sesamoid bone fatigue in Thoroughbred racehorses is associated with palmar osteochondral disease in the adjacent third metacarpal bone condyle"

### Supplementary Figures and Tables

*Elleana J. Thom,[1,3] *Soroush Irandoust PhD,[1,2] Corinne R. Henak PhD,[2-4] Peter Muir BVSc, MVetClinStud, PhD, DACVS, DECVS, FRCVS[1]

[1]Comparative Orthopaedic Research Laboratory, School of Veterinary Medicine, University of Wisconsin-Madison, Madison, WI 53706 USA

[2]Department of Mechanical Engineering, University of Wisconsin-Madison, Madison, WI 53706, USA

[3]Department of Biomedical Engineering, University of Wisconsin-Madison, Madison, WI 53706, USA

[4]Department of Orthopedics & Rehabilitation, University of Wisconsin-Madison, Madison, WI 53705, USA

*These authors contributed equally to the study

**Figure S1**. In the study 58 condyles and proximal sesamoid bones were analyzed from 22 Thoroughbred racehorses. In one limb, PSB comminuted impaired PSB segmentation for density estimation.

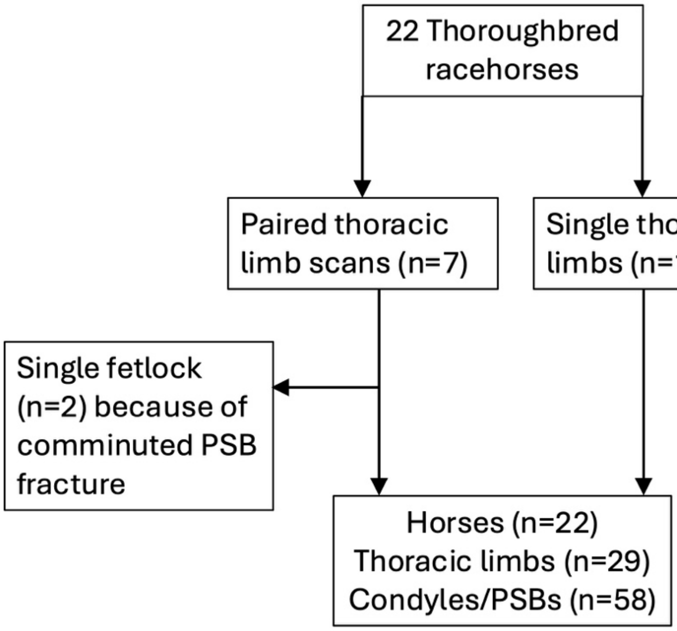

**Figure S2.** Case proximal sesamoid bones had significantly increased average HA-equivalent density that control bones (A) with a left-skewed distribution (B). Proximal sesamoid bones with mid-body or basilar stress fracture had qualitatively higher average HA-equivalent density than non-fractured bones, but the contrast was not significant (*P*=0.5) (C, D). Medial PSBs had significantly increased BMD than lateral PSBs (E, F). Shaded areas in the profile plots indicate the 95% confidence interval.

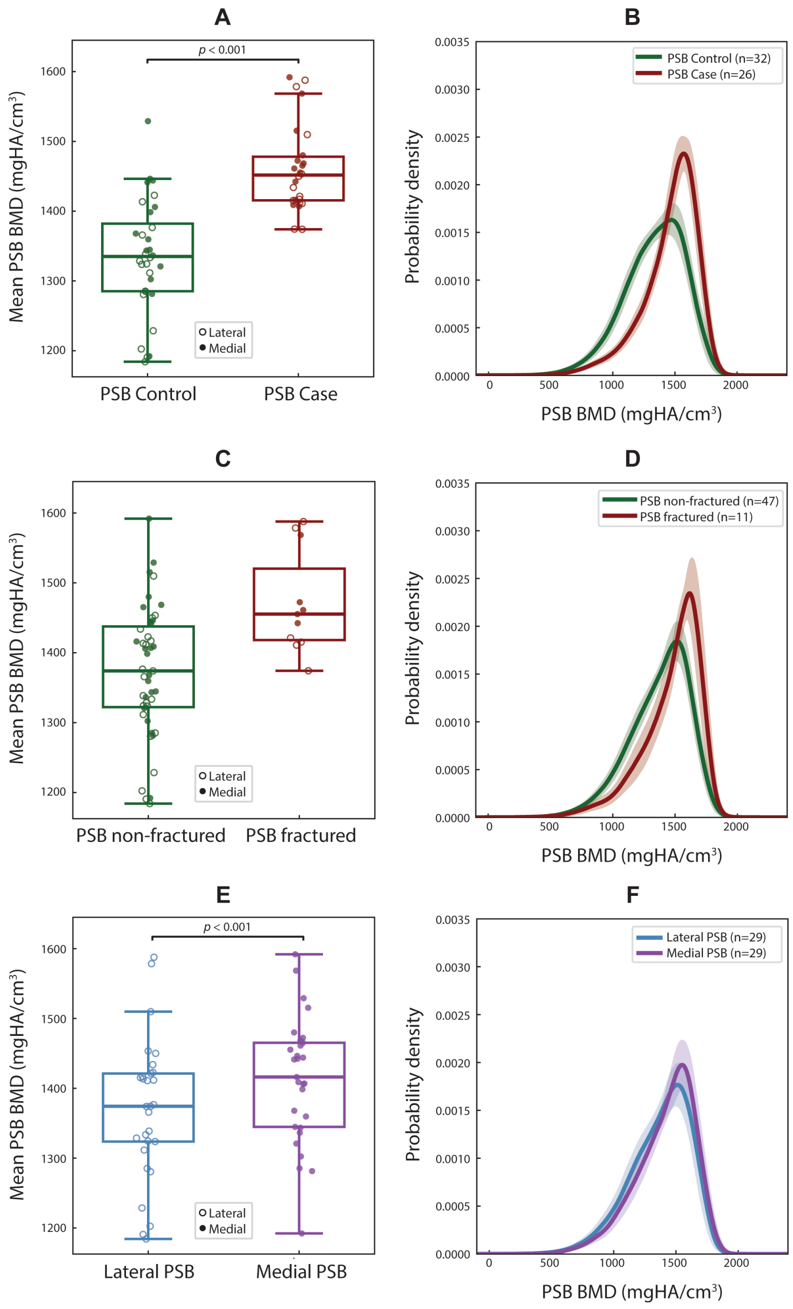

**Table S1**. Classification of structural changes associated with PSB fatigue

| | **Number of specimens** | **Proportion of 58 PSBs (%)** |
|---|---|---|
| **Mid-body transverse fracture with focal subchondral lysis and sclerosis** | 2 | 3.4 |
| **Mid-body transverse fracture with sclerosis** | 4 | 6.9 |
| **Basilar transverse fracture with focal lysis and sclerosis** | 1 | 1.7 |
| **Apical/basilar avulsion fracture with sclerosis** | 5 | 8.6 |
| **Subchondral focal lucency with sclerosis** | 4 | 6.9 |
| **Moderate to severe sclerosis** | 10 | 17.2 |
| **Structural change associated with bone fatigue** | 26 | 44.8 |
| **No structural change associated with bone fatigue** | 32 | 55.2 |

Note. Both mid-body transverse fractures with focal subchondral lysis and sclerosis affected the medial PSB with associated lateral mid-body PSB transverse fracture in a sclerotic bone and collapse of the fetlock.

**Table S2**. Association of PSB fatigue injury with POD fatigue injury

| | **POD case** | **POD Control** | **Total** |
|---|---|---|---|
| **PSB case** | 22 | 4 | 26 |
| **PSB Control** | 3 | 29 | 32 |
| **Total** | 25 | 33 | 58 |

Relative Risk = 0.03, Odds Ratio = 53.16, $P$>0.0001, Fisher Exact Test

| | **PSB case** | **PSB Control** | **Total** |
|---|---|---|---|
| **POD case** | 22 | 3 | 25 |
| **POD Control** | 4 | 29 | 33 |
| **Total** | 26 | 32 | 58 |

Relative Risk = 7.26, Odds Ratio = 53.16, $P$>0.0001, Fisher Exact Test

**Table S3**. Association of PSB fatigue injury with PSG fatigue injury

| | PSG case | PSG Control | Total |
|---|---|---|---|
| **PSB case** | 2 | 24 | 26 |
| **PSB Control** | 8 | 24 | 32 |
| **Total** | 10 | 48 | 58 |

Relative Risk = 0.31, Odds Ratio = 0.25, *P*=0.16, Fisher Exact Test

| | PSB case | PSB Control | Total |
|---|---|---|---|
| **PSG case** | 2 | 8 | 10 |
| **PSG Control** | 24 | 24 | 48 |
| **Total** | 26 | 32 | 58 |

Relative Risk = 0.40, Odds Ratio = 0.25, *P*=0.16, Fisher Exact Test

**Table S4**. Association of POD fatigue injury with PSG fatigue injury

| | POD case | POD Control | Total |
|---|---|---|---|
| **PSG case** | 3 | 7 | 10 |
| **PSG Control** | 22 | 26 | 48 |
| **Total** | 25 | 33 | 58 |

Relative Risk = 0.65, Odds Ratio = 0.51, *P*=0.49, Fisher Exact Test

| | PSG case | PSG Control | Total |
|---|---|---|---|
| **POD case** | 3 | 22 | 25 |
| **POD Control** | 7 | 26 | 33 |
| **Total** | 10 | 48 | 58 |

Relative Risk = 0.57, Odds Ratio = 0.51, *P*=0.49, Fisher Exact Test
